# Brainwide dopamine dynamics across sleep-wake transitions

**DOI:** 10.64898/2026.02.13.705774

**Authors:** Changwan Chen, Xun Tu, Lihui Lu, Cody Pham, Xiaofan Zhang, Rachel Su, Ella Li, Zihan Jin, Wenqing Cao, Yipching Yang, Matthew Kihiczak, Kristal Hui, Dana Darmohray

## Abstract

Dopaminergic signaling plays a critical role in regulating arousal and transitions between sleep states, yet brainwide dynamics underlying these transitions remain incompletely characterized. Here, we employed multi-site fiber photometry with the dopamine sensor GRAB-DA_2m_, combined with electroencephalogram (EEG)/electromyogram (EMG) recordings, to systematically assess regional dopamine (DA) dynamics across sleep-wake states in the medial prefrontal cortex (mPFC), striatal subregions, central amygdala (CeA), and midbrain nuclei in mice. We found that DA levels prominently increased in the mPFC, dorsolateral striatum (DLS), ventral tegmental area (VTA), substantia nigra pars compacta (SNc), and dorsal raphe nucleus (DRN) during transitions from non-rapid eye movement (NREM) sleep to wakefulness (WAKE), supporting their role in arousal. Conversely, DA decreased in the CeA and nucleus accumbens lateral parts (NAc-L) during these transitions. During transitions from NREM to rapid eye movement sleep (REM), DA elevations were observed in the CeA and middle accumbens subregion (NAc-M), rather than NAc-L, while other regions exhibited decreases. Cross-regional DA correlations revealed synchronized network activity across sleep-wake transitions. Optogenetic activation of VTA and DRN dopamine neurons induced robust DA release in cortical and subcortical regions, and chemogenetic activation promoted wakefulness selectively via VTA and DRN, but not SNc DA neurons. These results elucidate distinct brainwide DA dynamics across state transitions and highlight differential roles for DA signaling in modulating sleep-wake states.

## Introduction

Sleep–wake regulation emerges from coordinated activity across distributed neural circuits and neuromodulators systems^1–4^. Among these, dopamine plays a crucial role in arousal, and the gating of behavioral state transitions, with midbrain dopaminergic nuclei projecting broadly to cortical and subcortical targets^5–10^. Dopaminergic neurons in the VTA, SNc, and DRN innervate regions including the mPFC, striatal subregions, and the amygdala, positioning dopamine to influence both global brain states and region-specific computations^11–14^.

Prior studies have implicated dopamine in promoting wakefulness and shaping transitions between NREM and REM sleep^1,8,9,15,16^. However, most measurements of dopamine across sleep-wake transition have been constrained to single regions or limited sampling approaches, making it difficult to resolve how dopamine dynamics are coordinated across brainwide during state transitions. Moreover, the heterogeneity of dopaminergic signaling, together with mixed findings regarding its role in sleep regulation, leaves the organization of these signals into functional networks incompletely characterized.

To address these gaps, we asked: How do dopamine levels evolve across sleep–wake transitions in cortical, striatal, amygdalar, and midbrain regions? Are these dynamics synchronized across circuits, and do they reflect functional segregation across WAKE, NREM, and REM sleep? Which dopaminergic populations causally drive these patterns? In this study, we systematically characterized dopamine signaling using multi-site fiber photometry combined with EEG/EMG recordings. We identified brainwide spatiotemporal dynamics of dopamine activity during transitions between sleep and wake states.

## Results

### Distinct dynamic patterns of dopamine signaling during sleep-wake transitions

To investigate brainwide DA dynamics during different state transitions, we expressed the fluorescent DA sensor GRAB-DA_2m_^17^ and implanted optical fibers into 8 brain regions, including the mPFC, NAc-L, NAc-M, DLS, CeA, SNc, VTA, and DRN. Sleep–wake states were classified through simultaneous EEG and EMG recordings (Fig. 1A, B, Fig. S1).

**Figure 1.**
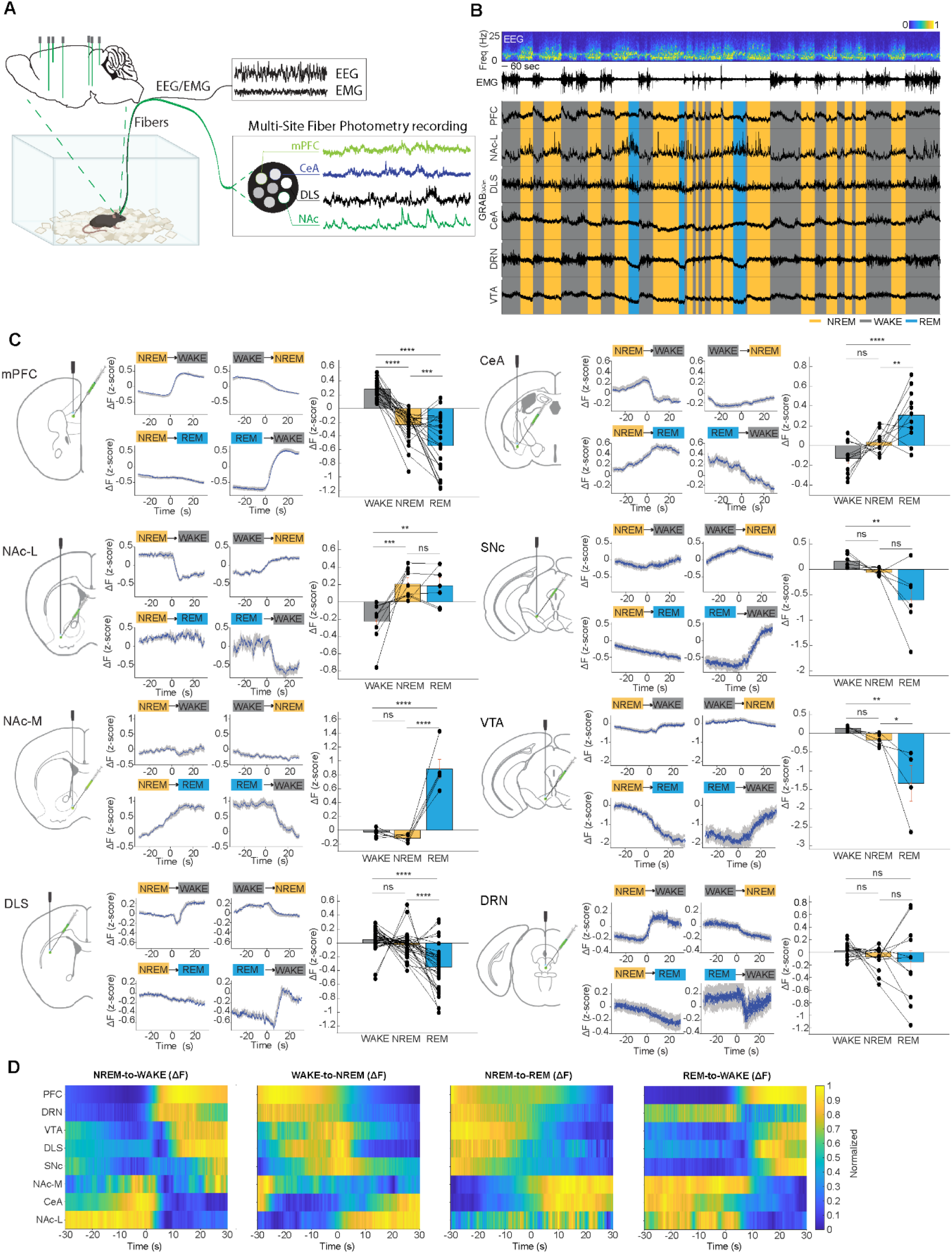
Distinct dynamic patterns of dopamine signaling during sleep-wake states. **(A)** Schematic illustration depicting the experimental design in which DA sensor was injected into different downstream targets and EEG/EMG were recorded. **(B)** An example multi-site fiber photometry recording session. Shown are the EEG spectrogram (normalized by the maximum of each session; Freq., frequency), EMG amplitude (Ampl.), brain states (color-coded), and DA signal traces. **(C)** Temporal changes of DA levels in different brain areas at each transition. For each subpanel: Left, schematic of the DA sensor injection and recording site; Middle, DA activity aligned to brain-state transitions (time zero indicates the transition point; shading represents ±s.e.m.); Right, summary of DA activity during WAKE, NREM, and REM, with each dot representing one animal. **(D)** Summary of DA activity (normalized to [0 1]) during 4 brain-state transitions across different brain areas. The data shown here is the average across mice.

In the mPFC, DA levels increased during transitions from NREM to WAKE and from REM to WAKE (Fig. 1B-D, Fig. S1). Conversely, DA levels decreased during WAKE-to-NREM and NREM-to-REM transitions. In contrast, DA dynamics in the CeA exhibited almost an opposite pattern, with the highest DA levels during REM sleep and the lowest during WAKE (Fig. 1B-D, Fig. S1).

Interestingly, DA dynamics in the striatum displayed distinct temporal patterns across its subregions. In the NAc-M, DA signaling exhibited a transient decrease during NREM-to-WAKE transitions, a robust decrease during REM-to-WAKE transitions, and a marked increase during NREM-to-REM transitions. In the NAc-L, DA levels decreased during NREM-to-WAKE and REM-to-WAKE transitions, increased during WAKE-to-NREM transitions, and exhibited no significant change during NREM-to-REM transitions. Conversely, DA levels in the DLS displayed an opposing trend, with the highest signals during WAKE and the lowest during REM. Notably, DA levels in the DLS also exhibited a transient dip during NREM-to-WAKE transitions (Fig. 1B-D, Fig. S1).

In brain regions containing dopamine neurons, the SNc and VTA shared similar DA dynamics, characterized by a pronounced increase during REM-to-WAKE transitions, a decrease during NREM-to-REM transitions, a slight increase during NREM-to-WAKE transitions, and a decrease during WAKE-to-NREM transitions. In contrast, DA dynamics in the DRN showed partially distinct dynamics, with a robust increase during NREM-to-WAKE transitions, a transient decrease during REM-to-WAKE transitions, and slight decreases during WAKE-to-NREM and NREM-to-REM transitions (Fig. 1B-D, Fig. S1).

Overall, DA levels in the mPFC, DLS, VTA, SNc, and DRN increased during NREM-to-WAKE transitions. During REM-to-WAKE transitions, DA levels in these regions also increased, except for the DRN. These results support the brainwide functional role of DA signaling in wakefulness. DA levels in NAc-L, NAc-M, and CeA decreased during both NREM-to-WAKE and REM-to-WAKE transitions, and increased during NREM-to-REM transitions, which is consistent with previous findings that DA signaling is elevated during NREM and/or REM sleep in these brain areas^15,18^.

### DA signaling displays distinct correlation patterns between regions during state transitions

To understand the relationships between different DA signaling during state transitions, we next computed cross-correlations between DA signals in different brain regions and analyzed their temporal dynamics across state transitions. During NREM-to-REM and REM-to-WAKE transitions, DA dynamic changes were generally more synchronized between regions than those during NREM-to-WAKE and WAKE-to-NREM transitions, with less correlation peaks- or troughs-shifting from the time lag *t* = 0 s (NREM-to-REM: |*t*| = 1.41 ± 0.48; REM-to-WAKE: |*t*| = 1.21 ± 0.33; NREM-to-WAKE: |*t*| = 3.84 ± 0.76; WAKE-to-NREM: |*t*| = 4.55 ± 1.01. Fig. 2A, 2C, Fig. S2A, 2C).

**Figure 2.**
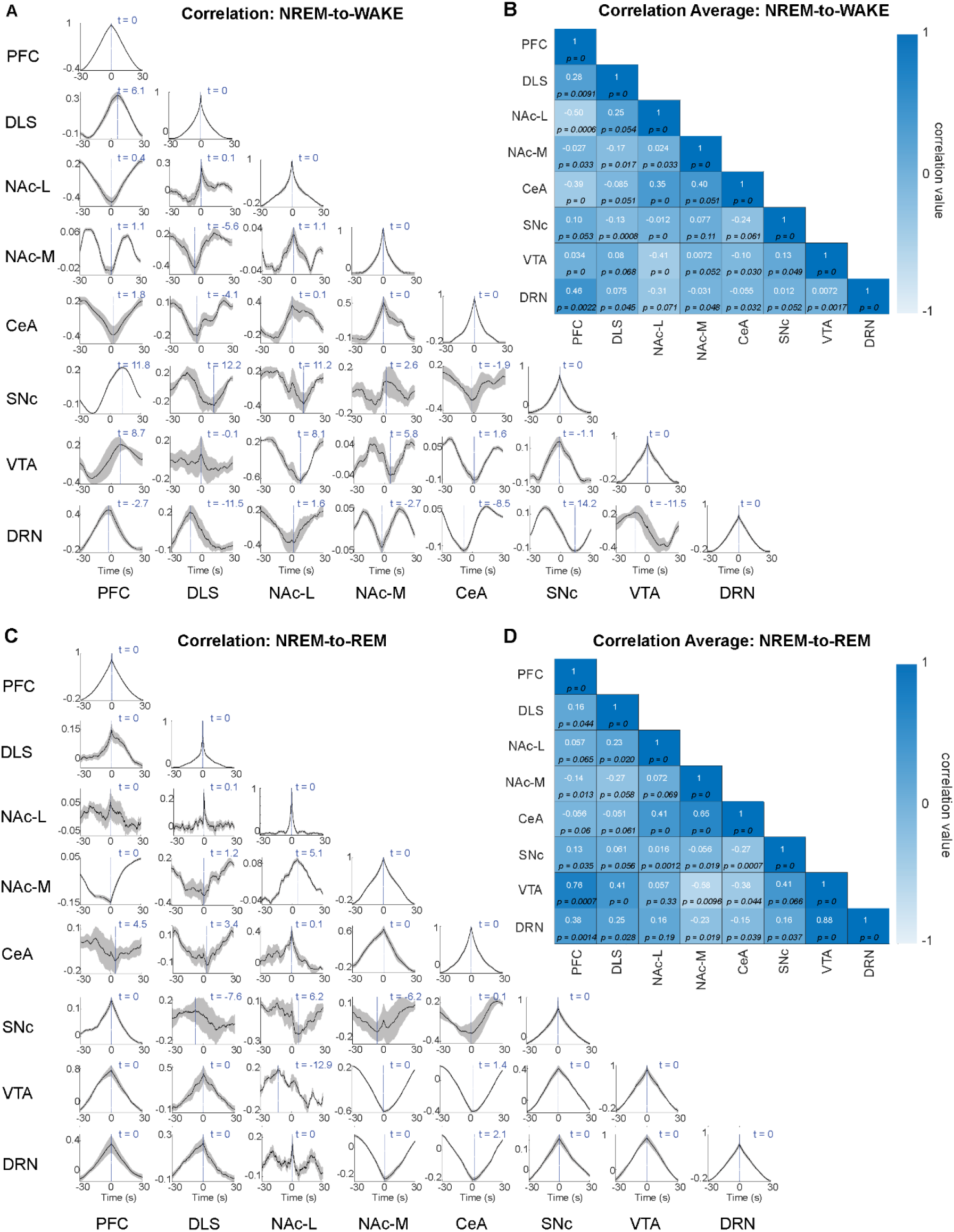
Correlation patterns of dopamine release during sleep-wake transitions. **(A)** Temporal correlation of DA signaling during NREM to WAKE transition states between different brain areas recorded simultaneously. **(B)** Correlation coefficient of the DA signals during NREM to WAKE transition states between regions. **(C)** Temporal correlation of DA signaling during NREM to REM transition states between different brain areas recorded simultaneously. **(D)** Correlation coefficient of the DA signals during NREM to REM transition states between regions.

Across all sleep-wake transition states, DA signaling in the CeA was highly correlated with that in the both NAc-L and NAc-M, as indicated by significant cross-correlation between regions (Fig. 2B, 2D, Fig. S2B, 2D), which also reflected in the temporary correlation dynamics, with high positive values of the correlation peaks at around *t* = 0 s (Fig. 2A, 2C, Fig. S2A, 2C). Similar correlation dynamics were also observed between PFC and DLS (Fig. 2, Fig. S2). DA signals in these two regions were positively correlated with DA signals in the SNc, VTA, and DRN, respectively, during NREM-to-REM transition and REM-to-WAKE transition. However, DA signals in the NAc-M and CeA are negatively correlated with DA signals in the SNc, VTA, and DRN during these transitions. Interestingly, the NAc-L and DLS had a strong correlation across all the transition states (Fig. 2, Fig. S2).

Furthermore, during NREM-to-REM transition, peak correlations in DA signals between the PFC and the SNc, VTA, and DRN occurred at *t* = 0 s, indicating a high degree of synchrony across these regions. In contrast, during the NREM-to-WAKE transition, DA activity in the DRN increased prior to that in the PFC, whereas DA signals in the SNc and VTA rose only after PFC activity increased. DA dynamics in the striatum regions exhibited regional heterogeneity by comparing the correlations with PFC DA signaling during both NREM-to-REM and NREM-to-WAKE transitions. All DA signaling in the NAc-L, NAc-M, and DLS changed after that in the PFC during NREM-to-WAKE transition, and were more synchronized at *t* = 0 s during NREM-to-REM transition. In the CeA, DA signaling changed later than that in the PFC across all transitions (Fig. 2A, 2C).

### Laser-evoked VTA, SNc, and DRN dopamine signaling exhibit distinct response patterns

To map the sources of dopaminergic inputs to key brain regions, including the mPFC, DLS, NAc, and CeA, we employed an integrated approach combining optogenetic activation of dopamine neurons with multi-site fiber photometry recording to monitor real-time dopamine activity. Targeted optogenetic stimulation was achieved using the red-shifted opsin ChrimsonR, enabling precise temporal control of DA neuronal activation while recording. Simultaneous EEG/EMG recordings were conducted to correlate dopamine dynamics with sleep-wake states^19^.

Activation of dopamine neurons in the VTA elicited robust dopamine responses in the mPFC, NAc, DLS, and CeA. Notably, the response amplitudes were larger in the NAc and DLS compared to the mPFC and CeA (dopamine responses in the mPFC, NAc, DLS, CeA: 2-s stimulation, mean peak amplitude *A* = 0.35, 2.04, 1.63, 0.17, respectively. 10-s stimulation: *A* = 0.36, 1.66, 1.59, 0.20 respectively. Fig. 3A-C). These findings align with the anatomical projections of VTA dopamine neurons^11,20–22^, which are denser in the striatum than in the mPFC or CeA^13^. Furthermore, Prolonged optogenetic stimulation was associated with more sustained dopamine responses, indicating a temporal dependence in the response dynamics.

**Figure 3.**
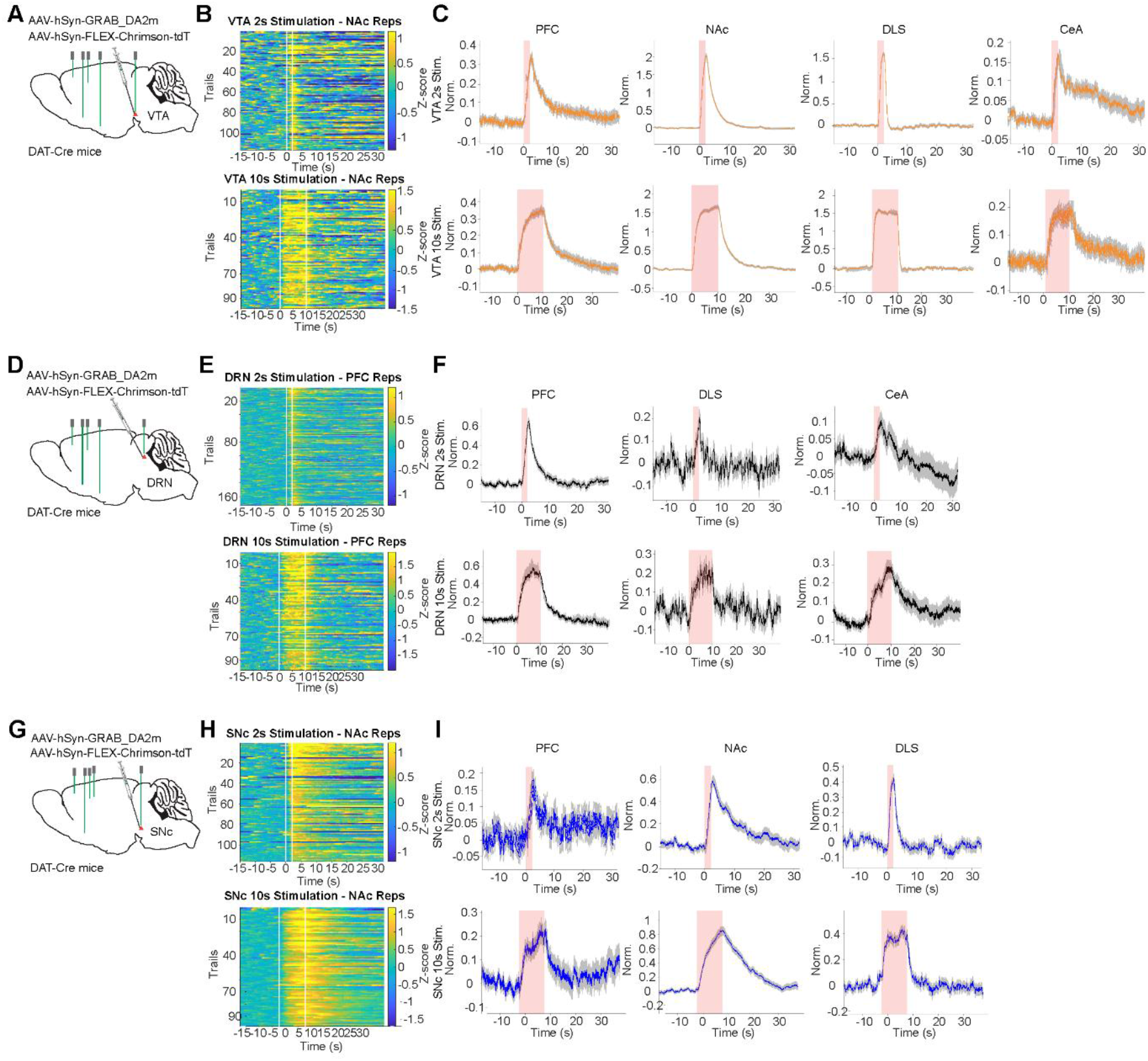
Laser-evoked VTA and DRN dopamine signaling exhibit distinct response patterns. **(A)** Schematic illustration depicting the experimental design in which ChrimsonR and GRAB-DA_2m_ vectors were injected into the VTA, and the DA sensor was injected into different downstream targets. **(B)** Heatmap of laser-evoked DA response in NAc in one session. Top, 2 s stimulation protocol; Bottom, 10 s stimulation protocol. **(C)** Laser-evoked DA sensor response in PFC, NAc, DLS, and CeA with the 2 s (averaged peak amplitude: *A* = 0.35, 2.04, 1.63, 0.17 respectively) or 10 s (averaged peak amplitude: *A* = 0.36, 1.66, 1.59, 0.20 respectively) stimulation in the VTA. **(D)** Schematic illustration depicting the experimental design in which ChrimsonR and GRAB-DA_2m_ vectors were injected into the DRN, and the DA sensor was injected into different downstream targets. **(E)** Heatmap of laser-evoked DA response in PFC in one session. Top, 2 s stimulation protocol; Bottom, 10 s stimulation protocol. **(F)** Laser-evoked DA sensor response in PFC, DLS, and CeA with the 2 s (averaged peak amplitude: *A* = 0.65, 0.45, 0.21, 0.10 respectively) or 10 s (averaged peak amplitude: *A* = 0.77, 0.70, 0.29, 0.29 respectively) stimulation in the DRN. **(G)** Schematic illustration depicting the experimental design in which ChrimsonR and GRAB-DA_2m_ vectors were injected into the SNc, and the DA sensor was injected into different downstream targets. **(H)** Heatmap of laser-evoked DA response in NAc in one session. Top, 2 s stimulation protocol; Bottom, 10 s stimulation protocol. **(I)** Laser-evoked DA sensor response in PFC, NAc, and DLS with the 2 s (averaged peak amplitude: *A* = 0.18, 0.57, 0.30, 0.43 respectively) or 10 s (averaged peak amplitude: *A* = 0.24, 0.86, 0.23, 0.44 respectively) stimulation in the SNc.

Optogenetic activation of DRN dopamine neurons produced region-specific increases in DA levels within the mPFC, DLS, and CeA (2-s stimulation: mean peak amplitude *A* = 0.65, 0.21, and 0.10; 10-s stimulation: *A* = 0.77, 0.29, and 0.29; Fig. 3D–F), in a pattern that mirrors previously described anatomical projections from the DRN to these target regions^12,13^.

Consistent with previously described projection patterns^23^, activation of SNc dopamine neurons evoked DA responses in the NAc, DLS, and mPFC, with the largest amplitudes observed in the NAc (2-s stimulation: mean peak amplitude *A* = 0.18, 0.57, and 0.43; 10-s stimulation: *A* = 0.24, 0.86, and 0.44; Fig. 3G–I).

These widespread responses suggest that dopamine neurons may exert a broad influence across multiple brain areas involved in regulating sleep-wake states and transitions between them.

### Activating VTA or DRN but not SNc dopamine neurons promotes wakefulness

To further assess the functional roles of midbrain dopamine source neurons, we expressed a chemogenetic vector *AAV-hSyn-FLEX-Gq* in VTA, SNc, or DRN dopamine neurons of *DAT-Cre* mice and implanted EEG/EMG electrodes to classify brain state changes. In line with prior studies showing that optogenetic activation of VTA dopamine neurons promotes wakefulness^9^, we found that CNO-induced activation significantly increased wakefulness while concomitantly reducing both NREM and REM sleep (Fig. 4A, B, G-H). Although both VTA and DRN dopamine neuron activation increased wakefulness and reduced NREM and REM sleep, the effect was more pronounced following VTA activation (Fig. 4C, D, G-H), consistent with prior reports on their roles in arousal regulation^8,9^. Activation of SNc dopamine neurons produced minimal changes in wakefulness and NREM sleep, suggesting a comparatively limited role in arousal regulation (Fig. 4E, F, G-H). Control mice lacking Gq expression showed no significant changes in brain states following CNO injection, confirming the specificity of the chemogenetic manipulation (Fig. S3).

**Figure 4.**
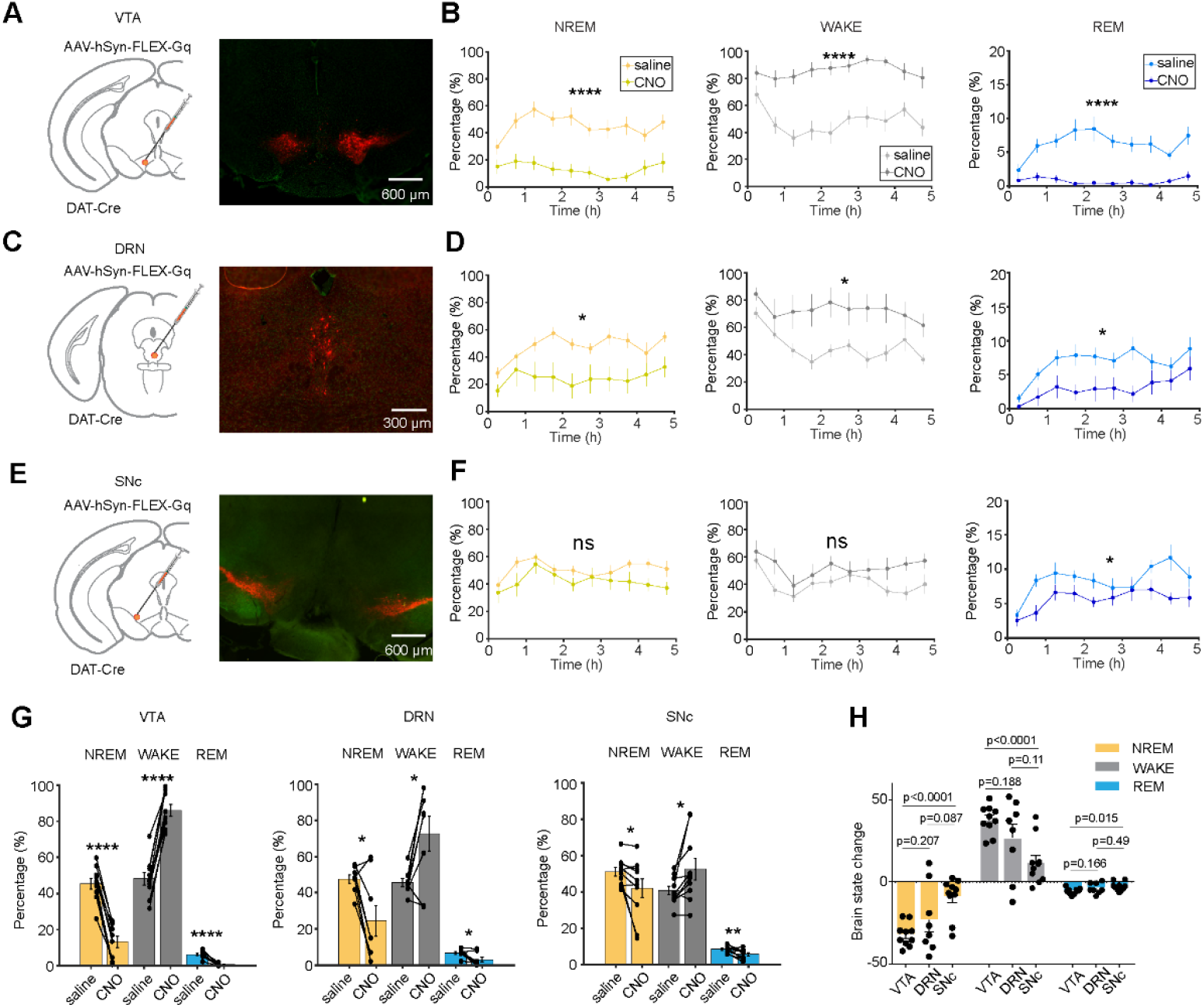
Activating VTA or DRN dopamine neurons promotes wakefulness. **(A)** Schematic illustration depicting the experimental design in which Gq vectors were injected into the VTA. The example image shows viral expression in the VTA. **(B)** Effect of VTA DA neurons activation on sleep and wake. Summary of the percentages of time in each brain state following CNO or saline injection in VTA. **(C)** Schematic illustration depicting the experimental design in which Gq vectors were injected into the DRN. The example image shows viral expression in the DRN. **(D)** Effect of DRN DA neurons activation on sleep and wake. Summary of the percentages of time in each brain state following CNO or saline injection in DRN. **(E)** Schematic illustration depicting the experimental design in which Gq vectors were injected into the SNc. The example image shows viral expression in the SNc. **(F)** Effect of SNc DA neurons activation on sleep and wake. Summary of the percentages of time in each brain state following CNO or saline injection in SNc. **(G)** Summary of average percentages of time in each brain state. **(H)** Changes in each brain state induced by chemogenetic activation (difference between CNO and saline injections, averaged across 5 h after injection) in mice with Gq expression in VTA, DRN, or SNc.

Collectively, these results reveal differential roles of midbrain dopamine neuron subpopulations in in arousal: VTA and DRN dopamine neurons strongly promote wakefulness, whereas SNc dopamine neurons exert comparatively limited influence (Fig. 4G-H).

## Discussion

Our study provides a comprehensive brainwide map of dopamine dynamics across sleep-wake transitions, revealing both region-specific and state-specific signatures of DA signaling. Notably, increases in DA levels during transitions from NREM sleep to WAKE were prominent in the mPFC, striatal regions, VTA, SNc, and DRN, consistent with established wake-promoting functions of these dopaminergic circuits^3,8,9,24,25^. However, distinct DA decreases in the CeA and NAc-L during transitions to WAKE, and DA elevations in these regions during NREM to REM transitions, suggest that DA plays specific roles in NREM and REM sleep regulation^15,26,27^, contributing to its initiation and maintenance through pathways distinct from those governing wakefulness.

The correlation analyses further support the presence of functional dopaminergic networks. Dopamine signals in amygdalar and nucleus accumbens subregions displayed highly synchronized dynamics, which were negatively correlated with cortical and midbrain DA fluctuations during state transitions. These patterns suggest a functional segregation of dopaminergic signaling— potentially driven by differences in propagation, integration, or local modulation—wherein some circuits are specialized for arousal and others are preferentially engaged during NREM and REM sleep^26–29^, which align with prior work implicating VTA, SNc, and DRN DA neurons in sleep-wake regulation^8,9,15,18^.

Furthermore, functional differences in DA signaling are likely attributable, at least in part, to regional variations in the kinetics of DA release. To the best of our knowledge, our study provides the first systematic mapping of DA signaling kinetics across key brain regions by selectively stimulating dopaminergic neurons in the SNc, VTA, or DRN. Specific DA responses are also consistent with previous studies that characterized DA signaling from the SNc to striatal subregions^24^ and from the VTA to striatal and amygdalar subregions^15,30,31^. Further studies are needed to determine whether specific DA signaling circuits are necessary and/or sufficient to regulate sleep-wake states.

Chemogenetic activation of VTA and DRN DA neurons strongly promoted wakefulness, consistent with previous studies using optogenetic activation of VTA^9^ and DRN DA neurons^8^, respectively. While activation of SNc DA neurons produced only mild effects on sleep-wake states, this may be attributed to the functional heterogeneity of DA neurons within the SNc. Three genetically defined dopamine neuron subtypes—characterized by expression of *Slc17a6, Calb1*, and *Anxa1*—have demonstrated significant differences in accelerations and decelerations responses, which may explain the distinct functional profiles of SNc DA neurons in sleep-wake regulation compared to those in the VTA and DRN^25^.

Taken together, our results delineate distinct brainwide functional patterns of DA activity during sleep-wake transitions and highlight the complexity of DA-mediated sleep-wake regulations. Future studies are needed to systematically probe how specific DA circuits govern the transitions between different behavioral states.

## Materials and Methods

### Animals

All procedures were performed in accordance with the protocol approved by the Animal Care and Use Committee at the University of California, Berkeley. Adult (>8 weeks old) male and female Dat-Cre (*Slc6a3*^*tm1(cre)Xz*^/J. RRID:IMSR_JAX:020080) mice were used for all experiments. Mice were kept on a 12:12 light:dark cycle (lights on at 7:00 a.m. and off at 7:00 p.m.) at constant room temperature and humidity with free access to food and water. After Surgeries (virus injections and implantation of EEG/EMG electrodes and optical fibers), mice were individually housed. Recording experiments were conducted 3-5 weeks after surgery.

### Surgeries

For all surgeries, anesthesia was induced with 5% isoflurane and maintained with 1.5% isoflurane on a stereotaxic frame. Buprenorphine (0.1 mg/kg) and meloxicam (10 mg/kg) were injected before surgery. Lidocaine (0.5%, 0.1 mL) was injected near the target incision site. The body temperature was kept stable throughout using a heating pad and a feedback thermistor. Eye ointment was applied to keep the eyes from drying. After sterilizing the skin with ethanol and betadine, an incision was made to the skin to expose the skull. Surgeries typically consisted of virus injections followed by EEG/EMG implantation and fiber implantation.

For virus injections, a craniotomy was drilled on the top of the target site, and AAVs were injected into the target regions using a Nanoject II (Drummond Scientific) with glass pipettes. The injection settings were set to a 300 nL injection volume/site at a rate of 23 nL/s with a 15-s interval between injections. pAAV9-hysn-GRAB_DA2m (titer: 2.4 × 10^13 gc/mL, from WZ Biosciences Inc and Addgene) and pAAV-hsyn-GRAB_DA-mut (titer: 2.7 × 10^13 gc/mL, from WZ Biosciences Inc and Addgene) was injected into target sites including:

The medial prefrontal cortex (mPFC): anteroposterior (AP) +1.94 mm, mediolateral (ML) +(/−)0.3 mm, dorsoventral (DV) −2.3–2.5 mm

The nucleus accumbens medial part (NAc-M): AP +1.50 mm, ML +(/−)0.55 mm,DV −4.6–4.8 mm

The nucleus accumbens lateral part (NAc-L): AP +1.42 mm, ML +(/−)1.25 mm, DV −4.6–4.8 mm The Dorsolateral Striatum (DLS): AP +0.98 mm, ML +(/−)2.3 mm, DV −3.0–3.2 mm

The central nucleus of the amygdala (CeA): AP −1.3 mm, ML +(/−)2.6 mm, DV −4.8–4.9 mm

The substantia nigra pars compacta (SNc): AP −3.08 mm, ML +/−1.4 mm, DV −4.1–4.3 mm (bilaterally)

The ventral tegmental area (VTA): AP −3.4 mm, ML +/−0.4 mm, DV −4.3–4.5 mm (bilaterally) The dorsal raphe nucleus (DRN): AP −4.1 mm, ML 0 mm, DV −3.0-3.2 mm

For optogenetic and chemogenetic manipulations, AAV9-hSyn-FLEX-ChrimsonR-tdT (titer: 4.5 × 10^12 gc/mL, from Addgene, or AAV-Ef1a-DIO-rsChRmine-oScarlet-Kv2.1 (titer: 4.8 x 10^12, from UNC Vector Core) and AAV of AAV9-hSyn-DIO-hM3Dq-mCherry (titer: 2× 10^13 gc/mL, from WZ Biosciences Inc) were injected into SNc, VTA, or DRN. 10 minutes after the virus injection, the injection pipette was slowly removed from the injection site.

For recording the electroencephalogram (EEG) and electromyogram (EMG), stainless-steel wires (0.002” diameter) were soldered to a 20-pin header (Minitek 127T series). EEG screw electrodes were implanted in both sides of the frontal lobe (± 1.5 mm ML, −1.5 mm AP). EMG stainless-steel wire electrodes (0.003” diameter) were inserted into both sides of the trapezius muscle. Reference screws were inserted into each side of the cerebellum. For fiber photometry and optogenetic stimulation, optical fiber implants (1.25-mm ferrule, 200-µm core) were held by a stereotactic fiber holder and slowly implanted into target sites. The EEG/EMG implant was secured by dental cement after fiber implantation.

mPFC: AP +1.94 mm, ML +(/−)0.3 mm, DV −2.5 mm (or angularly 30.38°)

NAc-M: AP +1.50 mm, ML +(/−)0.55 mm, DV −4. 8 mm (or angularly 15.2°)

NAc-L: AP +1.42 mm, ML +(/−)1.25 mm, DV −4. 8 mm

DLS: AP +0.98 mm, ML +(/−)2.3 mm, DV −3. 2 mm (or angularly 14.48°)

CeA: AP −1.3 mm, ML +(/−)2.6 mm, DV −4. 9 mm

SNc: AP −3.08 mm, ML +/−1.4 mm, DV −4.3 mm (bilaterally)

VTA: AP −3.4 mm, ML +/−0.4 mm, DV −4. 5 mm (bilaterally, one fiber was implanted angularly 21. 8°)

DRN: AP −4.1 mm, ML 0 mm, DV −3. 2 mm (or angularly 17.88°)

Fibers were secured with cyanoacrylate glue and dental cement before withdrawing the stereotactic fiber holder.

### Sleep recording

Sleep behavioral experiments were carried out in home cages placed in sound-attenuating boxes between 9:00 a.m. and 7:00 p.m. Mice were habituated in the home cages with EEG/EMG cables connected. Each session was recorded for 4–5 h. Each mouse was typically recorded for ≥ 2 sessions. EEG and EMG electrodes were connected to flexible recording cables by a mini-connector. The signals were recorded with a TDT RZ5 amplifier (Synapse software), filtered (0– 300 Hz) and digitized at 1,500 Hz. Spectral analysis was carried out using fast Fourier transform (FFT). For each 5-s epoch, the brain state was classified into NREM, REM or wake states on the basis of EEG and EMG data (wake: desynchronized EEG and high EMG activity; NREM: synchronized EEG with high-amplitude, low-frequency (0.5–4 Hz) activity and low EMG activity; REM: high power at theta frequencies (6–9 Hz) and low EMG activity). Consecutive epochs of the same state were combined into a single episode. The classification was performed semi-automatically using a custom-written graphical user interface (programmed in MATLAB 2022a, MathWorks). For the comparison of normalized EEG power spectra within each brain state, the EEG spectra were normalized to the total EEG power between 0 and 25 Hz.

### Optogenetic stimulation

To study the laser evoked responses of dopamine release or sleep behavior changes induced by activation of either VTA or DRN dopamine neurons, we applied laser stimulation (635 nm, 10-mW at fiber tip). For laser evoked responses of dopamine release, two stimulation protocols were applied: (1), 30 Hz (10 ms pulse) with 2 seconds epochs for 2h stimulations, the inter-stimulation-interval was 1 minute between epochs. (2), 30 Hz (10 ms pulse) with 10 seconds epochs for 2h stimulations, the inter-stimulation-interval was 1 minute between epochs. Mice were recorded for 1h as baseline, and recorded 1h more post-stimulation.

### Chemogenetic manipulation

For chemogenetic manipulation, saline (0.9% sodium chloride) or CNO (C0832, Sigma; dissolved in saline) was injected i.p. into mice expressing hM3Dq or mice without hM3Dq expression. The CNO dose was 1 mg kg−1 body weight for hM3Dq or control experiments. Each recording session started immediately after injection and lasted for 5 h. Each mouse was recorded for 6–8 sessions, and CNO was given randomly in half of the sessions, while saline was given in the other half. Data was averaged across all sessions.

### Fiber photometry recording

Fiber photometry was performed using a Multichannel Fiber Photometry System (R820, RWD Life Science), with 470 nm and 410 nm LEDs. The sampling rate for each channel was 30 frames/s. The system was synchronized with TDT sleep recording system by TTL pulses. For analysis of GRABDA2m signals, we fit the 470 and 410 nm channels to a bi-exponential decay function to approximate the slow photobleaching time course. The bi-exponential fit was subtracted from each channel, and the 410 nm channel was fit to the 470 nm channel using a least squares method. The final corrected 470-nm signal was z-scored for further analysis.

### Histology

To evaluate the location and strength of virus expression as well as the fiber position, mice were anesthetized with isoflurane before transcardial perfusion with 20 mL 1× PBS (Gibco) and 20 mL 4% paraformaldehyde (Electron Microscopy Sciences) in 1× PBS. For fiber localization, the mouse head with implants was fixed in 4% paraformaldehyde for >24h before separating the brain from the top of the skull. Then the brain was tripped out from the skull, and further fixed with 4% paraformaldehyde for 24h. The fixed brain was dehydrated in 20 mL of 30% sucrose PBS for 24 h, allowing time for it to sink to the bottom of a conical tube. The brain was then embedded in tissue freezing medium (General Data Company, Inc.) and frozen for at least half an hour. The brain was cryosectioned at 40 µm using a cryostat (Leica). Slides were then imaged on a fluorescence microscope (Keyence).

### Statistical analysis

All statistical analyses were performed using MATLAB v2022a (MathWorks) and Prism9 v4.0 (GraphPad). All statistical comparisons were conducted on animals. The number of mice and the level of significance are indicated in the figure legends. Two-group comparisons were performed using a t-test. Multiple group comparisons were performed using either two-way or one-way ANOVA, following multiple two-group tests with α correction, adjusting for multiple comparisons. All statistics are reported as the mean ± s.e.m.

## Acknowledgments

We sincerely thank Prof. Yang Dan for her invaluable support, including funding, laboratory space, equipment, and exceptional guidance and supervision throughout this work. We thank Dr. Daniel Silverman for providing some photometry analysis code. We also appreciate the administrative assistance provided by Dr. Hongfeng Gao. Additionally, we are grateful to all members of the Dan Lab for their insightful discussions and collaborative support.

## Author contributions

Conceptualization: C.C., X.T., and L.L. Methodology: C.C., X.T., L.L. Investigation: C.C., X.T., L.L., C.P., X.Z., R.S., E.L., Z.J., W.C., Y.Y., M.K., K.B.H., D.D., and D.S. Data analysis: C.C., X.T., L.L. Writing, review, and editing: C.C., X.T., and L.L.

## Data and code availability

Any data and code for this paper are available upon request.

---

**Supplemental Figure 1.**
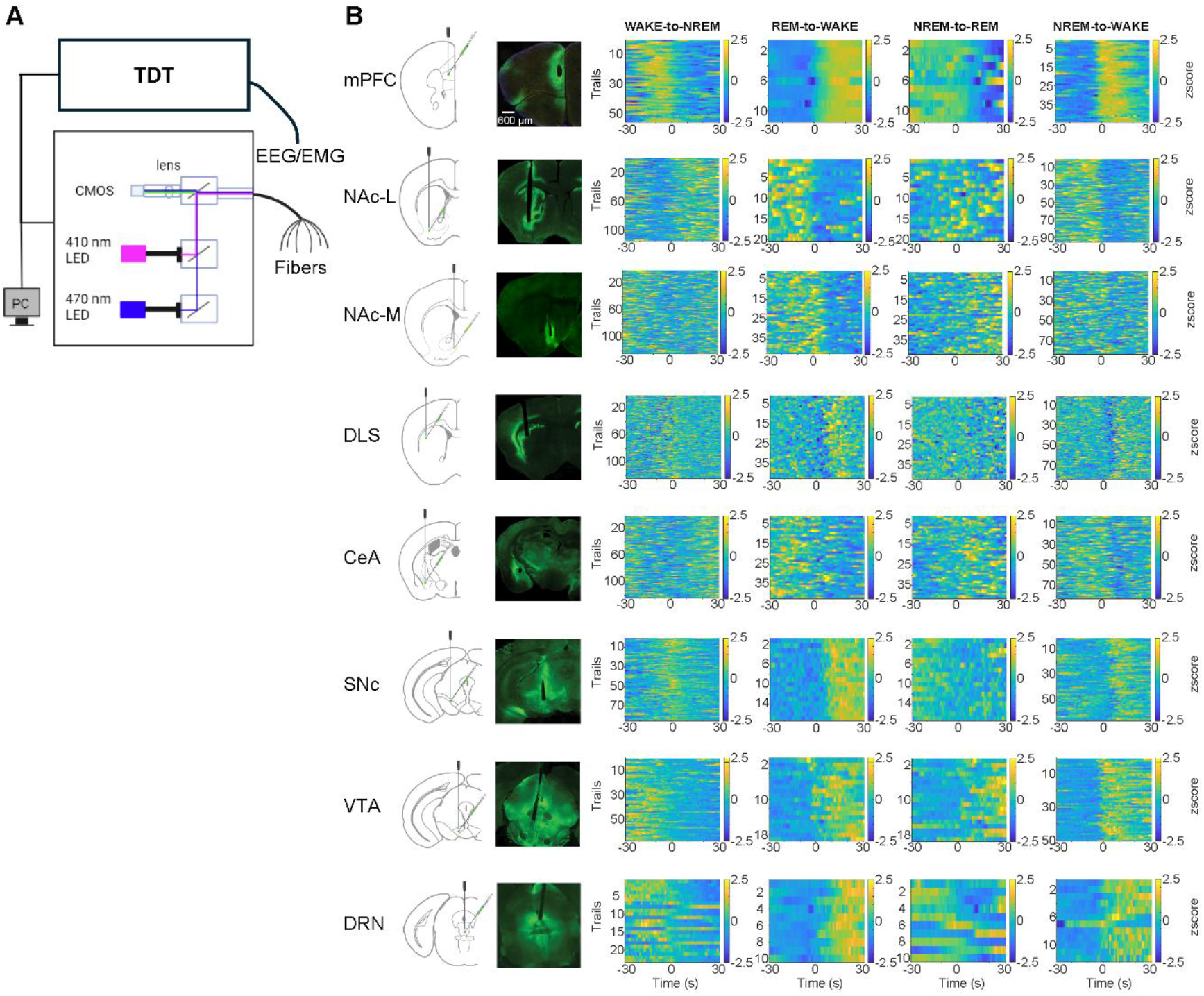
Dopamine dynamics across brain-state transitions in distinct brain regions. **(A)** Schematic of multi-site fiber photometry and EEG/EMG recording. TDT, Tucker-Davis Technologies. **(B)** Example of DA dynamics across different brain-state transitions. For each row: Left, Schematic of DA sensor injection; middle, example image of DA sensor expression; right, heatmap of DA signal across different transitions.

**Supplemental Figure 2.**
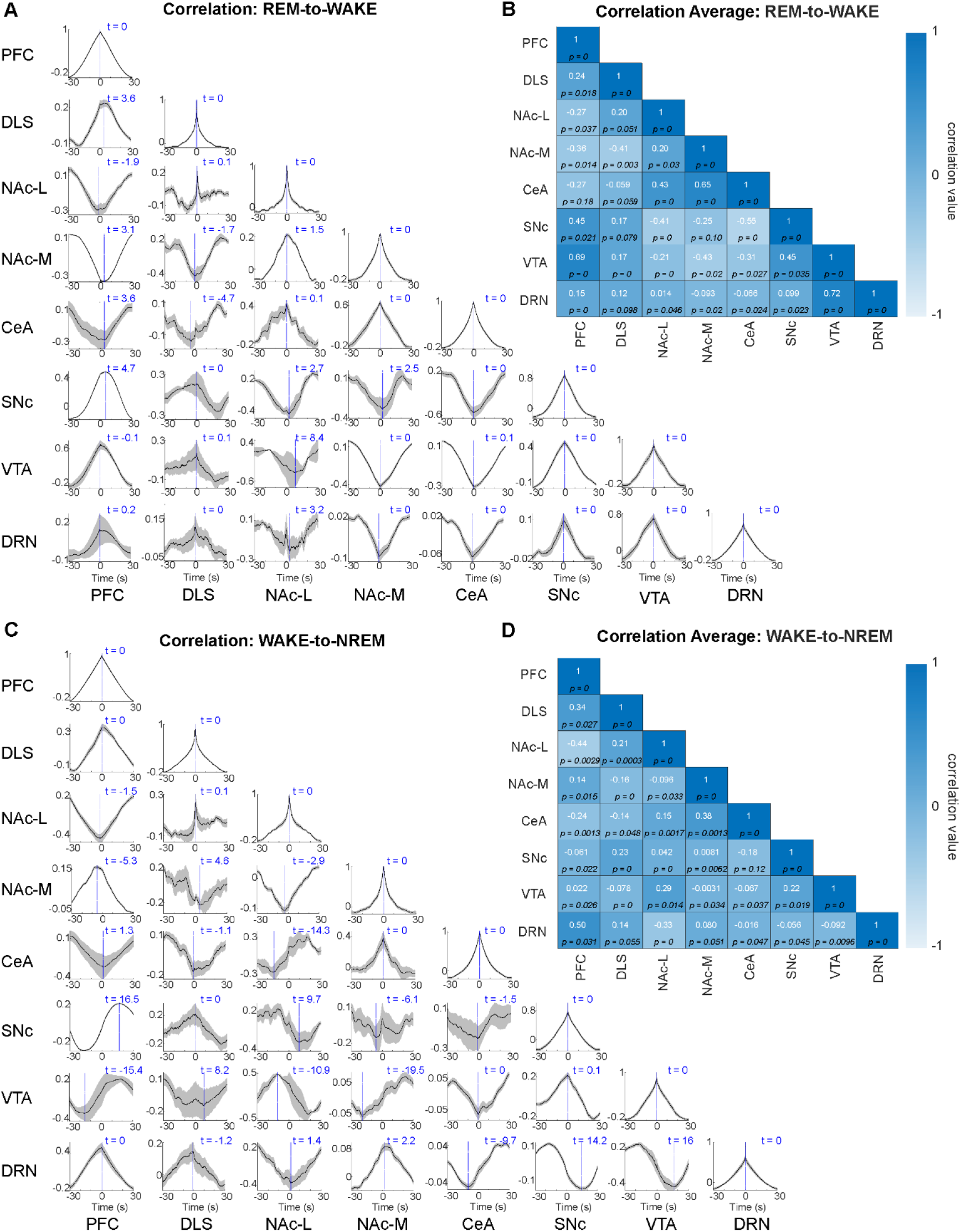
Correlation patterns of dopamine release during sleep-wake transitions. **(A)** Temporal correlation of DA signaling during REM to WAKE transition states between different brain areas recorded simultaneously. **(B)** Correlation coefficient of the DA signals during REM to WAKE transition states between regions. **(C)** Temporal correlation of DA signaling during WAKE to NREM transition states between different brain areas recorded simultaneously. **(D)** Correlation coefficient of the DA signals during WAKE to NREM transition states between regions.

**Supplemental Figure 3.**
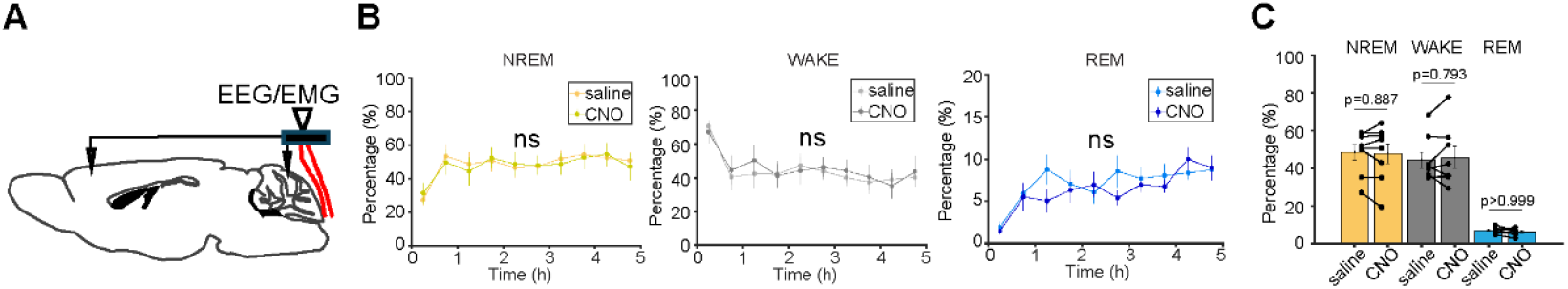
Control experiment for chemogenetic activation. **(A)** Schematic of EEG/EMG recording in mice without Gq vectors expression. **(B)** Summary of the percentages of time in each brain state following CNO or saline injection in DRN. **(C)** Summary of averaged percentages of time in each brain state.

**Supplemental Table.**

| Figure # | Conditions | n per group | Analysis | Factor | P value | P value summary |
| --- | --- | --- | --- | --- | --- | --- |
| 1C | PFC | 29 | One-way ANOVA with multiple comparisons | WAKE vs NREM | < 0.0001 | **** |
|  |  |  |  | WAKE vs REM | < 0.0001 | **** |
|  |  |  |  | NREM vs REM | 0.0006 | *** |
|  | NAc-L | 9 | One-way ANOVA with multiple comparisons | WAKE vs NREM | 0.0005 | *** |
|  |  |  |  | WAKE vs REM | 0.0019 | ** |
|  |  |  |  | NREM vs REM | 0.96 | ns |
|  | NAc-M | 5 | One-way ANOVA with multiple comparisons | WAKE vs NREM | 0.79 | ns |
|  |  |  |  | WAKE vs REM | < 0.0001 | **** |
|  |  |  |  | NREM vs REM | < 0.0001 | **** |
|  | DLS | 37 | One-way ANOVA with multiple comparisons | WAKE vs NREM | 0.25 | ns |
|  |  |  |  | WAKE vs REM | < 0.0001 | **** |
|  |  |  |  | NREM vs REM | < 0.0001 | **** |
|  | DMS | 4 | One-way ANOVA with multiple comparisons | WAKE vs NREM | 0.99 | ns |
|  |  |  |  | WAKE vs REM | 0.19 | ns |
|  |  |  |  | NREM vs REM | 0.15 | ns |
|  | CeA | 11 | One-way ANOVA with multiple comparisons | WAKE vs NREM | 0.094 | ns |
|  |  |  |  | WAKE vs REM | < 0.0001 | **** |
|  |  |  |  | NREM vs REM | 0.0048 | ** |
|  | SNc | 6 | One-way ANOVA with multiple comparisons | WAKE vs NREM | 0.6 | ns |
|  |  |  |  | WAKE vs REM | 0.0086 | ** |
|  |  |  |  | NREM vs REM | 0.059 | ns |
|  | VTA | 5 | One-way ANOVA with multiple comparisons | WAKE vs NREM | 0.64 | ns |
|  |  |  |  | WAKE vs REM | 0.0038 | ** |
|  |  |  |  | NREM vs REM | 0.017 | * |
|  | DRN | 14 | One-way ANOVA with multiple comparisons | WAKE vs NREM | 0.67 | ns |
|  |  |  |  | WAKE vs REM | 0.44 | ns |
|  |  |  |  | NREM vs REM | 0.91 | ns |
| 4B | NREM | 10, 10 | Two-way ANOVA | saline vs CNO | < 0.0001 | **** |
|  | WAKE | 10, 10 | Two-way ANOVA | saline vs CNO | < 0.0001 | **** |
|  | REM | 10, 10 | Two-way ANOVA | saline vs CNO | < 0.0001 | **** |
| 4D | NREM | 8, 8 | Two-way ANOVA | saline vs CNO | 0.0204 | * |
|  | WAKE | 8, 8 | Two-way ANOVA | saline vs CNO | 0.0168 | * |
|  | REM | 8, 8 | Two-way ANOVA | saline vs CNO | 0.0213 | * |
| 4F | NREM | 10, 10 | Two-way ANOVA | saline vs CNO | 0.1319 | ns |
|  | WAKE | 10, 10 | Two-way ANOVA | saline vs CNO | 0.0812 | ns |
|  | REM | 10, 10 | Two-way ANOVA | saline vs CNO | 0.0158 | * |
| 4G | NREM | 10, 10 | Paired t test | saline vs CNO | < 0.0001 | **** |
|  | WAKE | 10, 10 | Paired t test | saline vs CNO | < 0.0001 | **** |
|  | REM | 10, 10 | Paired t test | saline vs CNO | < 0.0001 | **** |
| 4H | NREM | 8, 8 | Paired t test | saline vs CNO | 0.0159 | * |
|  | WAKE | 8, 8 | Paired t test | saline vs CNO | 0.0147 | * |
|  | REM | 8, 8 | Paired t test | saline vs CNO | 0.0132 | * |
| 4I | NREM | 10, 10 | Paired t test | saline vs CNO | 0.0414 | * |
|  | WAKE | 10, 10 | Paired t test | saline vs CNO | 0.0254 | * |
|  | REM | 10, 10 | Paired t test | saline vs CNO | 0.0079 | ** |
| 4J | NREM | 10, 8 | Unpaired t test | VTA vs DRN | 0.2067 | ns |
|  | NREM | 10, 10 | Unpaired t test | VTA vs SNC | < 0.0001 | **** |
|  | NREM | 8, 10 | Unpaired t test | DR vs SNC | 0.0869 | ns |
|  | WAKE | 10, 8 | Unpaired t test | VTA vs DRN | 0.1884 | ns |
|  | WAKE | 10, 10 | Unpaired t test | VTA vs SNC | < 0.0001 | **** |
|  | WAKE | 8, 10 | Unpaired t test | DR vs SNC | 0.1098 | ns |
|  | REM | 10, 8 | Unpaired t test | VTA vs DRN | 0.1657 | ns |
|  | REM | 10, 10 | Unpaired t test | VTA vs SNC | 0.0146 | * |
|  | REM | 8, 10 | Unpaired t test | DR vs SNC | 0.4905 | ns |
| S3B | NREM | 8, 8 | Two-way ANOVA | saline vs CNO | 0.3672 | ns |
|  | WAKE | 8, 8 | Two-way ANOVA | saline vs CNO | 0.331 | ns |
|  | REM | 8, 8 | Two-way ANOVA | saline vs CNO | 0.3066 | ns |
| S3C | NREM | 8, 8 | Tukey's multiple comparisons test | saline vs CNO | 0.8868 | ns |
|  | WAKE | 8, 8 | Tukey's multiple comparisons test | saline vs CNO | 0.7928 | ns |
|  | REM | 8, 8 | Tukey's multiple comparisons test | saline vs CNO | > 0.9999 | ns |

